# Complete mitochondrial genome of golden silk producer *Antheraea assamensis* and its comparative analysis with other lepidopteran insects

**DOI:** 10.1101/110031

**Authors:** Deepika Singh, Debajyoti Kabiraj, Hasnahana Chetia, Pragya Sharma, Kartik Neog, Utpal Bora

## Abstract

Muga (*Antheraea assamensis*) is an economically important silkmoth endemic to North-eastern part of India and is the producer of the strongest known commercial silk. However, there is a scarcity of -omics data for understanding the organism at a molecular level. Our present study decodes the complete mitochondrial genome (mitogenome) of *A. assamensis* and presents comparative analysis with other lepidopterans. Mitogenome is a 15,272 bp long AT rich (~80.2%) molecule containing 37 genes (13 PCGs, 22 tRNAs, 2 rRNAs) and a 328 bp long control region. The overall mitogenome arrangement was similar to the other lepidopterans. Two PCGs *cox1* and *cox2* were found to have CGA and GTG as start codons respectively like some lepidopterans. Typical clover-leaf shaped secondary structures of tRNAs were found with a few exceptions such as unstable DHU and TΨC loop in *tRNA*^*Ser1*^ and *tRNA*^*Tyr*^; significant number of mismatches (35) spread over 19 tRNAs. The control region contained a six bp deletion atypical of other *Antheraea* species. Phylogenetic position was consistent with the traditional taxonomic classification of Saturniidae. The complete annotated mitogenome is available in GenBank (Accession No. KU379695). To the best of our knowledge, this is the first report on complete mitogenome of *A. assamensis*.

## Introduction

Mitochondria are known to be the descendants of α-proteobacterium endosymbionts retaining numerous bacterial features (1). Apart from being power house of the cell, they are essential for various cellular processes like fatty acid metabolism, apoptosis and aging (2). These functions are carried out by extra-nuclear genes encoded within the mitochondrial genome (mitogenome) along with many nuclear-encoded genes. These extra-nuclear genes include protein-coding genes (PCGs), transfer RNA (tRNA) and ribosomal rRNA (rRNA) genes present in the mitogenome which is mostly circular and self-replicative. It also consists of several non-coding regions with the longest being control (AT-rich) region comprising of several conserved regions and repeats. Mitogenome has several unique features like maternal inheritance, small genome size, faster evolution rate, low or absence of homologous recombination, evolutionary conserved gene products and richness in genetic polymorphism which makes it a potential marker for barcoding, phylogeography and phylogenetic studies (3). It plays a potential role in molecular evolutionary studies by elucidating evolutionary models and substitution rates/pattern that vary timely and across sequences. As compared to individual genes, whole mitogenomes are more informative phylogenetic models attributed to its multiple genome level features like gene position, content, secondary structures of RNA and control region.

Over the past few decades, animal mitogenomes, particularly insects (~80% of the sequenced arthropods), have become widely used and ideal systems for comparative genomics and molecular systematics (3, 4). Presently, mitogenome sequencing can be carried out at a surging rate through next generation sequencing (NGS) which provides a platform for multiplex sequencing of mitogenomes in a single run. Hence, it circumvents the limitations of Sanger sequencing with primer walking method which is both time consuming and costly. More than 300 insect Lepidopteran mitogenomes have been sequenced till now using both the above technologies as found in GenBank. The mitogenome ranges from 14-16 kilo basepairs in majority of the lepidopterans and consists of 37 genes (13 PCGs, 2 rRNA genes, 22 tRNA genes) along with a control region. The PCGs encode 2 subunits of ATPase (ATP6 and ATP8), 3 subunits of cytochrome c oxidase (COI, COII and COIII), 1 subunit of cytochrome B (CYTB) and 7 subunits of NADH dehydrogenase (ND1, ND2, ND3, ND4, ND4L, ND5 and ND6). These proteins are responsible for oxidative phosphorylation (OXPHOS) as they form essential mitochondrial membrane-associated protein complex systems (3). The control region plays a role in the replication and transcription of mitogenome. Several studies have demonstrated the utilities and potential of mitochondrial PCGs (*cox1* and *cox2*) as barcode markers in the order Lepidoptera (5). Few studies have also elucidated the comparative analysis among the domesticated and non-domesticated lepidopteran mitogenomes (6, 7). While the frequency of mitogenome sequencing of lepidopterans has increased; the evolutionary relationships among many family members of the same order have been seldom investigated.

Saturniidae is the most diverse family of wild silkmoths (giant moths, royal moths and emperor moths); most of them are unexplored and might have potential significance in sericulture area (8). As far as the current updates, mitogenome data of eleven species have been sequenced and made available in GenBank (https://www.ncbi.nlm.nih.gov/). However, individual gene sequences of various wild species are also available without their complete mitogenome sequences. *Antheraea assamensis,* muga silkmoth is semi-domesticated and one of the economically important moths of the same family. It is endemic to Assam and its adjoining hilly areas located in the North- eastern part of India. It is a multivoltine and polyphagous moth that primarily thrives on two host plants *Persea bombycina* and *Litsea monopetala* (9). Its silk has wide applications in textile industries and great potential as a biomaterial due to its unique biophysical properties like golden lustre, tenacity and UV radiation absorption (10). Like other *Antheraea* species, it produces reelable silk which is one of the most expensive silks of the world. However, its semi-domesticated nature and extraction of fibroin directly from cocoon fibers is a limitation in its extensive rearing and prospects of global applicability as a biomaterial. The whole genome of *A. assamensis* is not yet available; however, *de novo* transcriptome data from our laboratory is available in GenBank (Accession Number- SRX1293136, SRX1293137 and SRX1293138).

In the present study, we report the whole mitogenome sequence of *A. assamensis* using NGS and comparative analysis of its sequences and genome architectures with that of the other lepidopterans belonging to various levels of scientific classification. The comparative study was based on several characteristics such as genome arrangement, PCGs, tRNAs, rRNAs, nucleotide composition, codon usage, evolutionary rates, gene divergence, conserved regions in control region, etc. Furthermore, phylogenetic trees inferred using datasets like concatenated nucleotide sequences of 13 PCGs and whole mitogenome sequences were analyzed to elucidate the relationships among lepidopteran insects. This study will thus facilitate better understanding of comparative and evolutionary biology of *A. assamensis* with the other lepidopteran insects.

## Results and Discussion

### Sample processing, sequencing and assembly

The total DNA extracted from fifth instar larvae of *A. assamensis* was checked for the integrity, quantity and purity and was found to be optimal (105 ng/µl by NanoDrop spectrophotometer and 234 ng/µl by Qubit fluorometer). The extracted DNA was then used for the enrichment of mitochondrial DNA through NEBNext Microbiome DNA Enrichment Kit. The nuclear DNA was settled down through the action of magnetic beads on methylated bases and clear supernatant containing mitochondrial DNA was isolated for library preparation. The prepared library profile showed that size of mitogenome fragments were in the range of 180 to 880 bp. However, total insert size distribution was from 300 to 1000 bp as it involved adapters (~120 bp) along with mitogenome fragments. In addition to the proper distribution of fragments, their concentrations (~27.2 ng/µl) were also found to be adequate for sequencing [Supplementary Figure S1]. The sequencing was carried out by paired-end sequencing technology with read lengths of 2 × 150 bp in Illumina NextSeq 500 sequencer. The sequencing resulted in 32,12,599 total number of raw reads, out of which around 28,69,153 high quality reads were processed for scaffold preparation. These reads were used for mitogenome scaffold preparation through *de novo* assembly of sequenced contigs using SPAdes-3.5.0, SSPACE and CAP3 program. Finally, a scaffold of 15,272 bp length was obtained which represents the whole mitogenome sequence of *A. assamensis*.

### Genome annotation, visualization and comparative analysis

The mitogenome sequence (obtained after sequencing) was annotated using MITOS web server where the tRNA and rRNA sequences were completely annotated. The PCGs were annotated with ORF Finder and NCBI-BLAST; and annotation of the control region was carried out with NCBI-BLAST, which successfully determined their exact location in the mitogenome. The annotated mitochondrial genome of *A. assamensis* seemed to be closed and circular structure comprising of 37 genes (13 PCGs, 22 tRNAs and 2 rRNAs) along with a non-coding control region over its 15,272 bp length. 24 genes were found to be encoded by the major coding strand or J-strand (+) and the remaining 13 genes by the minor coding strand or N-strand (-). The whole mito-map constructed using BRIG and its characteristics like gene location and arrangement are depicted in the Fig. 1. The full annotated mitogenome sequence and SRA data of *A. assamensis* were submitted to NCBI GenBank under the accession numbers KU379695 and SRR3948351 respectively.

**Figure 1.**
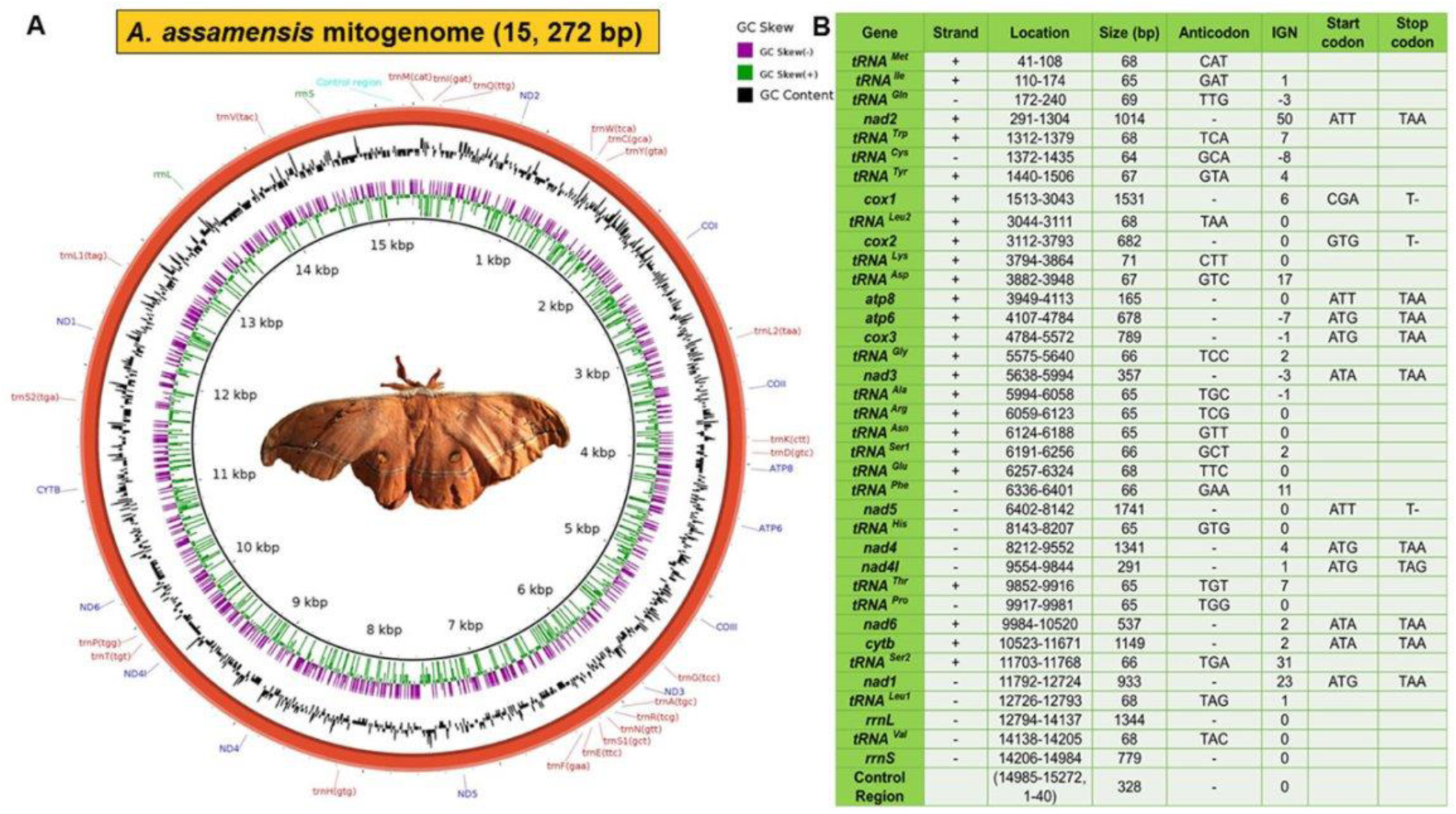
*A. assamensis* mitochondrial genome (A) Circular map showing gene content and rearrangement (B) Characteristic features of the mitogenome. Abbreviations: *cox1 cox2 and cox3*: cytochrome oxidase subunit I, II and III respectively, *cytb*: cytochrome B, *atp6* and *atp8*: subunits6 and 8 of F0 ATPase, *nad1-6*: components of NADH dehydrogenase. *tRNAs* are represented by *trn* followed by the IUPAC-IUB single letter amino acid codes e.g. *trnM, trnI* and *trnQ* denote *tRNA*^*Met*^, *tRNA*^*Ile*^ and *tRNA*^*Gln*^ respectively. The nucleotides in the bracket represent anticodons of *tRNAs*. IGN represents (+) values as intergenic nucleotides and (-) values as overlapping regions.

### Comparative mitogenome analysis

The mitogenome length of *A. assamensis* followed the characteristic size of most of the insects (14 to 20 kb) comprising of 37 genes and a control region (11, 12). This mitogenome size variation may be attributed to variation in non-coding regions especially the control region that shows great differences in length as well as pattern. It collectively leads to higher degree of gene rearrangements; however, PCGs are known to remain quite stable in the mitogenome. In order to study the conservation pattern; gene content, genome organization, structure, rearrangement, sequence similarity, etc. in the mitogenome of *A. assamensis* was compared with other lepidopterans belonging to various levels of scientific classification (Table 1).

**Table 1.**
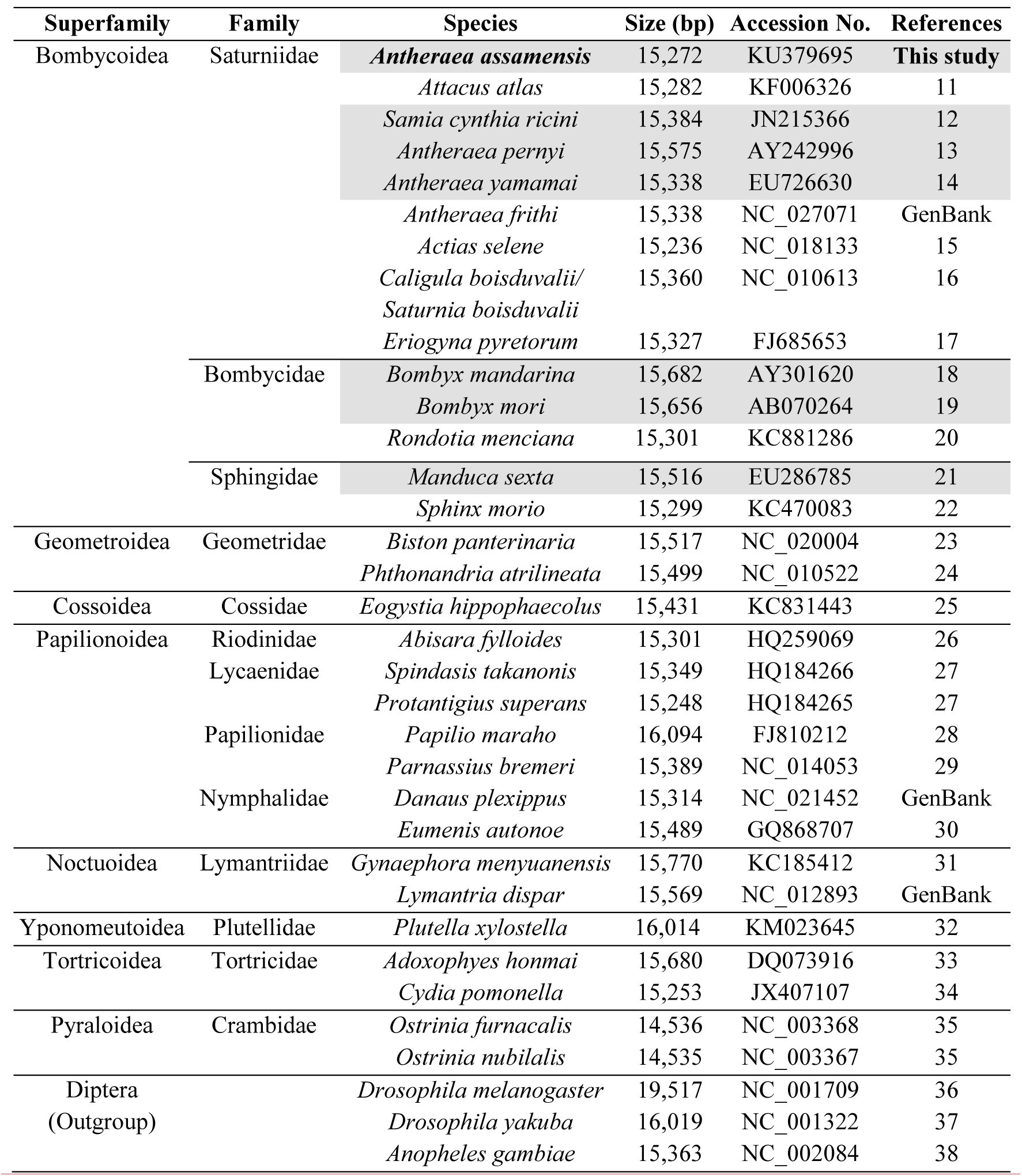
List of the organisms having sequenced mitogenome, belonging to Lepidoptera and Diptera (outgroup) order. (Highlighted ones are the organisms of Bombycoidea superfamily used for comparative mitogenome analysis with respect to *A. assamensis*).

The existence of a typical gene content i.e. 37 genes (13 PCGs, 22tRNAs, 2rRNAs) and a control region was evident in *A. assamensis* mitogenome as observed in other sequenced mitogenomes of lepidopteran insects (including Bombycoidea). The arrangement of mitochondrial genes was found to be in the order identical to the selected Saturniid and Sphingiid insects (*S. ricini, A. pernyi, A. yamamai* and *M. sexta*) while differing from Bombycids- *B. mori* and *B. mandarina*. Three typical rearrangements were observed in the mitogenome of *A. assamensis* similar to most of the other lepidopteran insects (23, 32, 33) as shown in Fig. 2. These were (i) *tRNA*^*Met-Ile-Gln*^ (*trnM/trnI/trnQ* or *MIQ)* cluster, (ii) *tRNA*^*Lys-Asp*^ cluster and (iii) *tRNA*^*Ala-Arg-Asn-Ser1-Glu-Phe*^ cluster. A typical *MIQ* cluster rearrangement located between the control region and *nad2* gene was found to be identical to Bombycoids and other insects of ditrysian lineages of lepidoptera (3, 12, 25). This arrangement differed from the ancestral pattern *tRNA*^*Gln-Ile-Met*^*(trnI/trnQ/trnM* or *IQM)* as reported in two ghost moths (Fig. 2) belonging to Hepialidae family - a non-ditrysian lineage (Lepidoptera: Exoporia: Hepialoidea) (39). Studies reveal that mitogenome rearrangement is multifactorial which is caused by various gene movements like inversion, transposition, inverse transposition, tandem duplication, etc. Also, multi-gene tRNA blocks are supposed to be hot spots for these rearrangements which seem to play a role in elucidating evolutionary mechanisms in insects (40). The *MIQ* cluster arrangement of *A. assamensis* exhibited the translocation of *trnM* which can be observed frequently in the evolutionary mechanism of insect mitogenome. This observation suggests that *A. assamensis* along with other lepidopteran insects have evolved with a typical gene arrangement after splitting from its stem lineage during the course of time. Further, sequence similarity performed among selected Bombycoids showed that *A. assamensis* shared highest similarity with *Antheraea* species (*A. yamamai* - 92.1%, *A. pernyi* - 91.7%) followed by the other Bombycoids like *S. ricini* (88.9%), *M. sexta* (85.5%), *B. mori* (83.4%) and *B. mandarina* (82.9%) indicating a significant level of homology among Bombycoidea superfamily than the other super families of Lepidoptera order.

**Figure 2.**
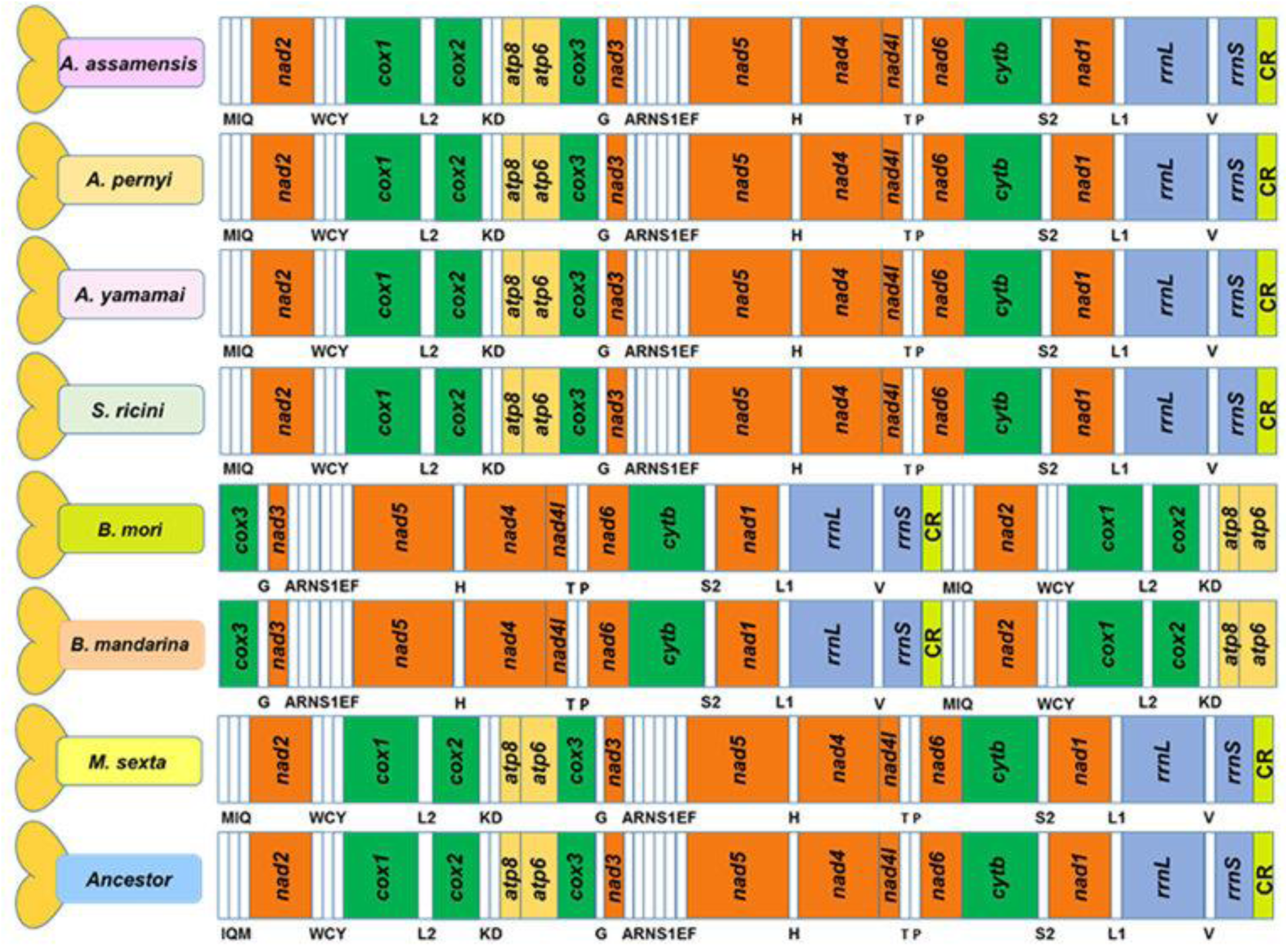
Arrangement of mitogenome in *A. assamensis* with respect to the related Bombycoid species. The arrangement of *Thitarodes renzhiensis* here denotes the ancestral type.

### Comparative analysis among protein coding genes (PCGs)

The mitogenome of *A. assamensis* comprised of a total of 13 PCGs which spanned 11,208 bp constituting around 73.38% of the total mitogenome. The total size of PCGs and their individual sizes were found to be lying within the range of other Bombycoid species (Supplementary Table S1). Similar to the other Bombycoid species, PCGs of *A. assamensis* were distributed over both the strands of double stranded mitogenome. Majority of genes (9 genes) such as *nad2, cox1, cox2, atp8, atp6, cox3, nad3, nad6* and *cytb* were encoded by J-strand and the remaining 4 genes (*nad5, nad4, nad4L* and *nad1*) by N-strand.

The PCG sequence analysis of *A. assamensis* showed ATN as the start codon for all the genes similar to the Bombycoid insects (12, 13, 14, 19, 21); specifically ATT, ATA and ATG were the starting codons for three, three and five genes respectively (Fig. 1B). Interestingly, start codons of *cox1* and *cox2* genes in *A. assamensis* were found to deviate from the canonical ATN initiation codon pattern. Similarly, *A. pernyi* (13), *A. yamamai* (14), *C. boisduvalii* (16), *B. mandarina* (18), *B. mori* (19) and *M. sexta* (21) were also reported to diverge from the pattern. In addition to the gene sequence similarity analysis, comparative amino acid sequence alignment of the reported lepidopteran insects and transcript data analysis of *A. assamensis* revealed that CGA and GTG were the initiation codons for both *cox1* and *cox2* genes of *A. assamensis* respectively. CGA was also found to be start codon for *cox1* gene in *Drosophila melanogaster* (41) and *Maruca vitrata* (42) through gene expression profiling. This was also reported in some of the members of Lepidoptera such as *A. atlas* (11), *E. pyretorum* (17), *M. sexta* (21), *A. honmai* (33), etc. On the other hand, GTG was found to be the start codon for *cox2* gene in *S. ricini* (12)*, C. boisduvalii* (16) and *E. pyretorum* (17).

Further, three genes *cox1, cox2* and *nad5* showed incomplete termination codon (only T) instead of canonical termination codon pattern TAA or TAG (Fig. 1B). The silkworm *A. pernyi* also showed incomplete termination codon in *cox1*, *cox2* and *nad5* genes (13). The existence of incomplete stop codon has been observed as a common phenomenon of mitochondrial genes of metazoans like lepidopterans in order to minimize intergenic spacers (IGS) and overlapping sequences (OS) (13, 19, 21, 29, 37). Several studies reveal that it could be a representative of recognition site for an endonuclease that truncates the polycistronic pre-mRNA. These incomplete codons are rectified by the post-transcriptional polyadenylation to yield functional stop codon with TAA termini (21).

In the next comparison study, PCG genes of *A. assamensis* were compared with seven different organisms belonging to various levels of scientific classification in Insecta class at nucleotide level and corresponding amino acid level in order to determine the conserved, variable and phylogenetically informative sites (Supplementary Table S2). Various species of *Antheraea* showed 84-90% conserved sites at nucleotide level while at amino acid level; it was comparatively higher (86-98%). However, *atp8* gene exhibited lower values (76.8%) at amino acid level which suggest that small variation in nucleotide has resulted in huge changes in amino acid composition. Similarly, various comparative analyses were carried out for the organisms of different genus, family, superfamily and order. The conserved sites were found to decrease with higher hierarchical organisms unlike variable and phylogenetically informative sites which were higher in number. Among all PCGs, *cox1* and *cox2* genes showed maximal conservation both at nucleotide and amino acid levels across the organisms of various classification levels.

Further, the pattern of nucleotide variation among the PCGs of *A. assamensis* with each family of Bombycoidea superfamily- *A. pernyi* (species level-Saturniidae), *B. mori* (family level-Bombycidae) and *M. sexta* (family level-Sphingidae) was studied. This pattern was found to vary between major and minor strands. While higher T-C transitions were observed in the PCGs encoded by the major strand, minor strand PCGs showed more A-G transitions when *A. assamensis* and *A. pernyi* were compared. These results were consistent with the earlier reported studies however, total number of transversions (178 No.s) was found to be higher than transitions (144) in the minor strand (7, 19). Similarly, between *A. assamensis* and *B. mori*, T-C transitions (299 T-C & 68 A-G over 5695 identical pairs) were found to be abundant on the major strand while A-G (155 A-G & 35 T-C over 3612 identical pairs) on the minor strand. In addition, the total number of transitions and transversions were found to be highest in the major strand between *A. assamensis* and *B. mori* followed by *M. sexta* and *A. pernyi*. This indicates high sequence divergence between *A. assamensis* and *B. mori* (Bombycidae) than with its closely related family members (Saturniidae) in Bombycoidea superfamily.

In order to determine the evolutionary rates, PCGs of *A. assamensis* were compared with seven different organisms belonging to various hierarchy levels in Insecta class. The average rate of synonymous substitutions (Ks) and average rate of non-synonymous substitutions (Ka) along with their ratios (Ka/Ks) were calculated for all 13 PCGs. The ratio Ka/Ks less than, greater than and equal to 1 indicates that genes are under negative (purifying) selection, positive (adaptative) selection and neutral evolution respectively (43). Among 13 PCGs, *atp8* gene encoding ATPase subunit 8 exhibited 1.5 ratio with reference to that of *D. melanogaster* indicating positive/ relaxed selection acting on this gene (Fig. 3). This shows that mutation in *atp8* of *A. assamensis* was due to a driving force to re-organize its structure. These changes may be due to the requirement of energy for the production and secretion of silk fibers which are not observed in *D. melanogaster*. It depicts that functional change in an organism needs commitment mutation in responsible genes in order to make survival of the fittest. Notably, the ratio for remaining PCGs of *A. assamensis* with all organisms was found to be less than one which suggests that mutation was against the requirement and hence mutation was replaced by synonymous nucleotides. However, Ka/Ks ratio in *atp8* with reference to *B. mandarina, B. mori* and *M. sexta* was found to be significantly higher. These organism falls in different family with respect to *A. assamensis* and hence significant variation in *atp8* gene may be expected due to certain changes in the organism kinetics. As *S. ricini, A. yamamai, A. pernyi* belong to the same family of *A. assamensis*, significant Ka/Ks ratio was not observed across the PCGs. This indicated that all the protein coding genes evolved under strong purifying (negative) selection. Overall comparison of PCGs exhibited that mutation in ATP8 subunit was essential in insects to yield several strains with different kinetics as other PCGs showed lower ratios. The lowest evolutionary rates were observed for *cox1* gene indicating that it is least susceptible to variation in protein sequence and hence shows potential as a barcode for evolutionary studies in silkworms (particularly Bombycoid species). In addition, other genes like *cox2, cox3* and *cytb* with slightly low evolutionary rates next to *cox1* can also serve as barcode markers. Furthermore, we studied whether the GC content has any significant effect on the Ka/Ks ratio and found a negative correlation between them (Supplementary Figure S2A). This indicates that change in GC content may result in the variation in nucleotide substitution pattern or evolutionary pattern among the PCGs.

**Figure 3.**
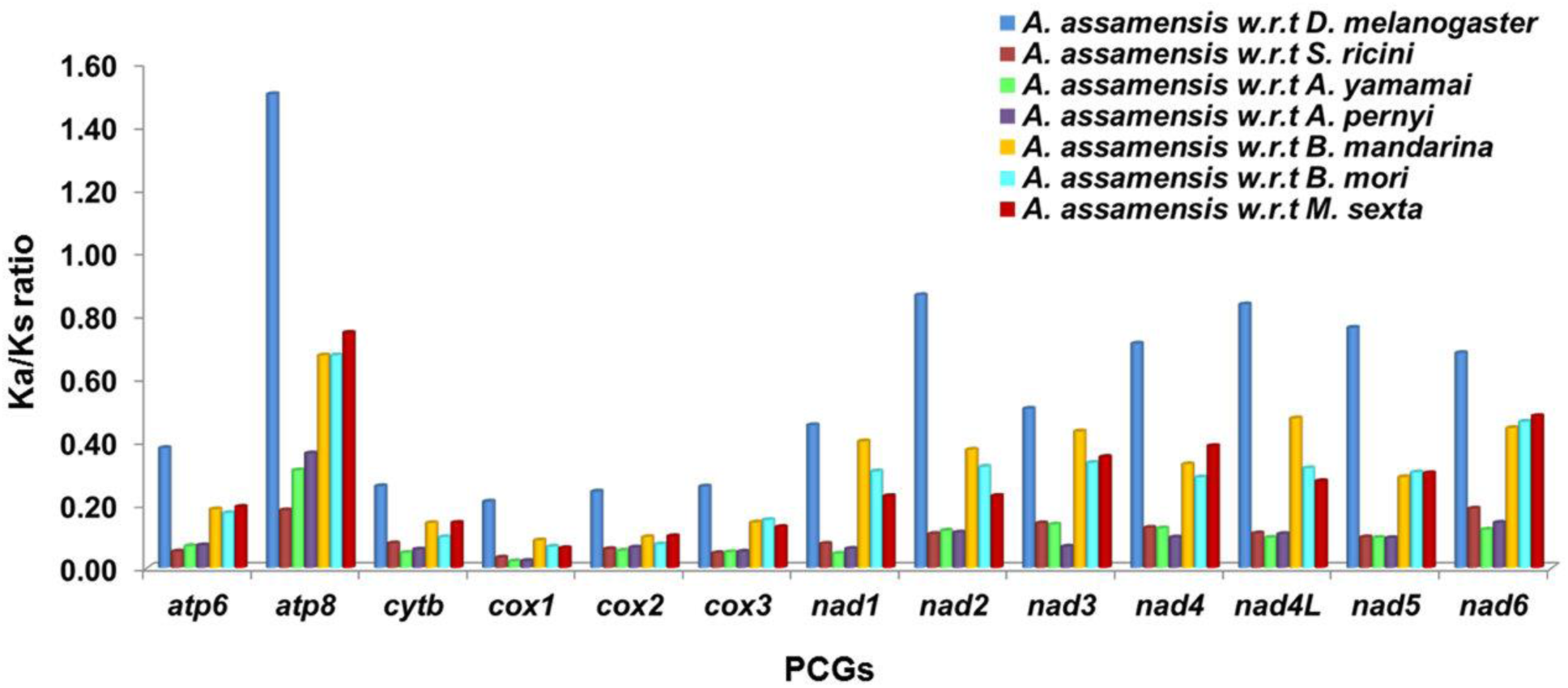
Evolutionary rates (Ka/Ks) of individual protein coding genes (PCGs) of *A. assamensis* in comparison with seven different organisms belonging to various hierarchy levels in Insecta class. w.r.t here denotes with reference to–.

### Comparative analysis among transfer RNAs (tRNAs)

Twenty two tRNA genes with a total length of 1465 bp were found to be present in the mitogenome of *A. assamensis*. The individual length of each gene varied from 64 to71 bp as reported for other lepidopterans (Supplementary Table S1). These tRNA genes were distributed over J and N strands of the mitogenome; 15 genes were encoded by J strand itself (Fig. 1B).

The secondary structures of tRNAs of *A. assamensis* exhibited a typical clover-leaf structure similar to other species of Lepidoptera order (see Supplementary Figure S3). This depicts that arrangement of all the tRNA genes is mostly conserved among insects. However, some variation in the structures of *A. assamensis* was observed such as aminoacyl acceptor stem of *tRNA*^*Met*^, TΨC loop of *tRNA*^*Ile*^, *tRNA*^*Cys*^, *tRNA*^*Tyr*^, *tRNA*^*Arg*^, *tRNA*^*Asn*^, *tRNA*^*Glu*^, *tRNA*^*Phe*^, *tRNA*^*Trp*^, *tRNA*^*Ser2*^, dihydrouridine (DHU) arm of *tRNA*^*Ser1*^, etc. The lack of stable DHU arm and TΨC loop has also been reported in lepidopterans such as *S. ricini, A. yamamai, A. pernyi, B. mori, B. mandarina*, *M. sexta*, *E. pyretorum, A. fylloides*, *B. panterinaria, E. autonoe,* etc. (12, 13, 14, 17, 19). These structural variations might be attributed to the variation in number of base pairs responsible for the formation of aminoacyl acceptor (AA) stem, DHU arm, TΨC arm and anticodon (AC) stem-loop (Supplementary Figure S4). Among various structural parts of tRNAs, AA stem of *A. assamensis* was found to be more consistent in size across all the tRNAs and in other lepidopterans as observed from comparative analysis. AC stem, AC loop and DHU stem also showed consistency in the number of base pairs across lepidopterans, however, varied with the type of tRNAs. In addition, anticodons of tRNAs were found to be identical to the selected Bombycoids (Supplementary Table S3) and majority of other lepidopteran insects (14, 16, 18, 23). On the contrary, TѰC stem, TѰC loop and DHU loop exhibited randomness in the number of base pairs across the organisms and tRNAs. For instance, TѰC loop was absent in *tRNA*^*Tyr*^ of *A. assamensis* while *tRNA*^*Trp*^ of *A. assamensis* and *tRNA*^*Tyr*^ of *B. mori* had 7 bp and 5 bp loops respectively. The number of base pairs of AA stem, AC stem, AC loop and DHU stem were highly consistent. This was probably to avoid dysfunctioning of tRNAs actively participating in protein synthesis.

Further, a total of 35 mismatched base pairs were identified in tRNAs of *A. assamensis* which were scattered over the AA stem, TΨC stem, AC and DHU regions of 19 tRNAs. 28 mismatches were observed in G-U combinations while 6 and 1 mismatches were found to be in U-U and G-A combinations respectively (Supplementary Table S3). These mismatches lead to changes in tRNA structures; for instance two U-U mismatches in *tRNA*^*Ser2*^ resulted in the formation of an extra loop in AC stem.

From the comparative analysis, we found that the number of total mismatches was higher in *A. assamensis* (35 No.s) than the selected Bombycoids like *S. ricini* (32 No.s), *B. mori* (29 No.s), *M. sexta* (25 No.s), etc. This number of mismatch was higher than many Saturniids like *E. pyretorum* (24 No.s), *S. cynthia* (28 No.s), etc. Similarly, mismatches such as G-U, U-U, G-A, A-A, A-C and C-U have been previously reported for many lepidopterans like *E. autonoe* (26 No.s), *A. fylloides* (27 No.s), etc. (26, 30). Among various mismatches, G-U mismatches were found be the highest (28 No.s) in *A. assamensis* than the selected Bombycoids and other lepidopterans e.g. *A. atlas* (25 No.s), *A. pernyi* and *E. autonoe* (20 No.s). The detailed comparative mismatches found in various stem-loop structures of different tRNAs of Bombycoid members are given in Supplementary Table S3. The non-canonical pairing might occur due to the insertion/deletion of nucleotides or nucleotide substitution of bases within the tRNA genes as a form of ancient insertional editing. The presence of high number of mismatches indicates higher nucleotide substitution in the tRNA sequences of *A. assamensis* as compared to the selected Bombycoids which may result in genome structure and sequence evolution. These mismatches are likely to be corrected by RNA editing mechanism as observed in other arthropods (44).

### Comparative analysis among ribosomal RNAs (rRNAs)

*A. assamensis* comprised of two genes for rRNA coded by (–) strand of the mitogenome. *rrnL* (lrRNA) is located between *tRNA*^*Leu1*^ and *tRNA*^*Val*^ genes, whereas *rrnS* (srRNA) is accommodated between *tRNA*^*Val*^ and control region as seen in other lepidopterans. The lengths of *rrnL* and *rrnS* were 1344 bp and 779 bp respectively similar to many reported lepidopteran insects (26, 31). The secondary structures of both rRNAs were predicted using Mfold and depicted in Supplementary Figure S5. The sequence homology of two rRNA genes of *A. assamensis* in comparison to selected Bombycoids revealed that the sequence identity of *rrnS* gene (84 to 95 %) was higher than that of *rrnL* (80 to 89 %). The divergence in rRNA genes (*rrnL* and *rrnS*) also exhibited low variability sites which suggest that these are highly conserved sites and may have potential application in molecular systematics.

### Comparative analysis among control or A+T region

The control region is the longest non-coding region of *A. assamensis* mitogenome and with a length of 328 bp, is similar to other Bombycoids. This region is located between *rrnS* and *MIQ* cluster. The comparative sequence homology studies of control region displayed higher similarity with its close relatives such as *A. yamamai* (89.2%) and *A. pernyi* (82.7%) than other selected Bombycoids (~68%). Based on the sequence alignment, some conserved structures and variable tandem repeats were observed in the control region of *A. assamensis* (Fig. 4). A motif ‘ATAGA’, poly T stretch microsatellite (TA)_9_ repeat sequence, poly-A region and 6 bp deletion were the major findings of study. These sequences are known to be important for gene regulation and serve as recognition site for replication initiation of minor or light strand. The poly A tail has been proposed to be required for RNA maturation and serve as a sequence for controlling transcription or replication initiation in insects (45). The microsatellites (T/A) also known as simple sequence repeats (SSR) are known to be used as molecular markers due to their abundance and highly polymorphic nature (12). Similarly, a microsatellite (AT)n element is a well conserved site and can be used for any conservative studies (13, 14). In our comparative analysis too we found highly conserved yet distinct consensus sequences in the control region. Thus this region has high potential as a phylogenetic marker at lower taxonomic levels.

**Figure 4.**
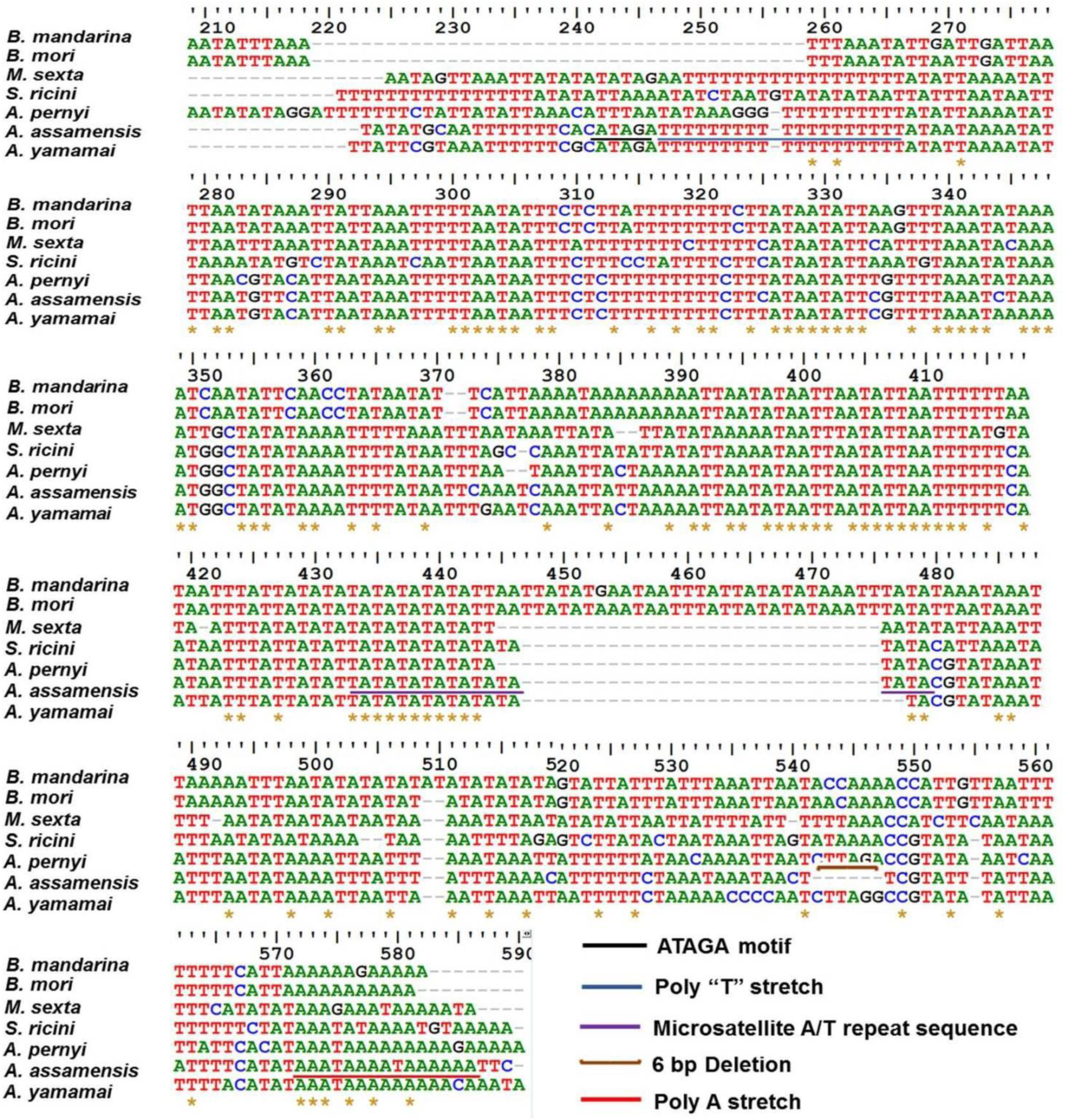
Multiple sequence analysis of control region of *A. assamensis* with selected Bombycoid species displaying conserved structures.

### Comparative analysis among overlapping sequence (OS) and intergenic spacer (IGS) regions

*A. assamensis* was found to have six OS with a total of 23 bp length however; individual sequence length was in the range of 1-8 bp as reported for many other lepidopterans (11, 15, 20, 31). Similarly, 17 IGS regions of total 171 bp length were found in *A. assamensis* mitogenome with a size range of 1 to 50 bp (Supplementary Table S4). The occurrence of OS and IGS is quite common in lepidopteran mitogenomes (12, 13, 14, 19). These regions may vary in length and location in order to reduce mitogenome size as a result of evolutionary pressure. A conserved overlapping junction (ATGATAA) was also found between *atp8* and *atp6* as reported for most of the lepidopteran mitogenomes (26, 30, 33).

*tRNA*^*Gln*^*-nad2* was found to be the largest spacer in *A. assamensis* (50 bp) which was slightly shorter than *A. yamamai* (53 bp), *S. cynthia* (54 bp), etc. Similarly, it has been detected as the largest in many other lepidopterans like *A. atlas, A. selene, C. boisduvali, E. pyretorum, P. bremeri* and *A. honmai* (11, 15, 16, 17, 29, 33). Other spacers such as *cytb-tRNA*^*Ser2*^ (31 bp), *tRNA*^*Ser2*^*-nad1* (23 bp), *tRNA*^*Lys*^*- tRNA*^*Asp*^ (17 bp), etc. also existed in *A. assamensis* (Fig. 1B). A conserved ‘ATACTAA’ motif was detected when IGS located between *tRNA*^*Ser2*^ and *nad1* was compared with selected Bombycoid species. The *cytb-tRNA*^*Ser2*^ spacer serves as a binding site for the mtTERM, a transcription termination peptide while *tRNA*^*Ser2*^*-nad1* is implicated in mitochondrial transcription termination where the ‘ATACTAA’ motif is essential as a recognition site for mtTERM (12, 14, 21).

Similar to some of the lepidopteran insects, the IGS in *A. assamensis* also displayed sequence homology with their adjacent genes. For instance, *tRNA*^*Gln*^-*nad2* spacer showed 68% similarity with its adjacent gene *nad2* in accordance with literature (14, 18, 21). The sequence homology of this spacer with its adjacent gene is due to the partial duplication of *nad2* gene as reported earlier and might serve as another origin of replication (12, 16, 21). This suggests that *A. assamensis* along with these organisms may have undergone rapid sequence divergence attributed to the non-coding nature of IGS (14). Apart from the adjacent genes, IGS in our organism showed sequence homology with their distant genes in the mitogenome as reported earlier in *S. ricini* (12). For example, spacer between *tRNA*^*Lys*^ and *tRNA*^*Asp*^ showed homology with *rrnS* (78%) and *nad5* (72%) genes in *S. ricini* while the same region in *A. assamensis* showed similarity with *rrnS* (82%) and *nad5* (76%) along with *atp8* gene (82%). Similar results have been found in the spacers *cytb-tRNA*^*Ser2*^ and *tRNA*^*Ser2*^*-nad1.* These findings suggest that occurrence of such homologous spacer sequences might be as a result of gene duplication and degeneration.

### Comparative analysis with respect to nucleotide composition, skewness and codon usage

The nucleotide composition in mitogenome of *A. assamensis* was analysed in terms of A+T content, AT and GC skewness. The nucleotide composition of the whole mito-genome of *A. assamensis* was found to be 40.8% thymine (T), 12.1% cytosine (C), 39.4% adenine (A) and 7.7% guanine (G). Thus with an A+T content of 80.2%, the muga mitogenome is biased towards A+T nucleotides like other lepidopterans ranging from *E. hippophaecolus* (78.4 %) to *S. takanonis* (82.3%) (25, 27). The control region exhibited maximal A+T content of ~91.2% followed by rRNAs (84.3%), tRNAs (80.8%) and PCGs (78%). The variation of A+T content with specific areas of mitogenome was found to be lying within the range of lepidopteran insects. Comparative AT content in the individual PCGs showed variation in the AT content where *atp8, nad4L* and *nad6* genes had the highest AT content while *cox1* and *cox3* genes had the lowest as detected in many Bombycoid species. This pattern was also identified in the other lepidopterans like *E. pyretorum, P. atrilineata, G. menyuanensis,* etc. (17, 24, 31). A close analysis of the third codon position in all 13 PCGs of *A. assamensis* exhibited AT biasness which was 93% than the first (73%) and second (70%) codon positions as similar to the other Bombycoids (12, 13, 17). This may be attributed to more relaxed constraints on A+T content at 3^rd^ position than at 1^st^ and 2^nd^ position due to the phenomena of degeneracy in genetic code.

Nucleotide skewness was determined to measure the relative number of Gs to Cs and As to Ts. The AT skewness in *A. assamensis* mitogenome was found to be negative (−0.02) which indicates the occurrence of more Ts than As and lies within the range for Saturniid moths (12, 13, 14). This value differed from the selected Bombycidae species having positive AT skew values e.g. 0.06 in *B. mori, B. mandarina*, etc. (18, 19). Similarly, mitogenome GC skewness also exhibited negative value (−0.22) as observed in many Bombycoids such as *M. sexta* (−0.18), *S. ricini* (−0.23). The negative values of these organisms indicate that they are rich in pyrimidines (T and C) throughout the mitogenome. The GC skewness varied with major and minor strands of PCGs, however no significant change was observed in AT skewness. The strand asymmetry is common in insects and has been reported in all the lepidopterans. These skewness patterns were also observed in rRNAs, tRNAs and the control region and are similar to the other sequenced lepidopteran insects (Supplementary Table S5).

Nucleotide biasness was also reflected in the codon usage of PCGs that determines the frequency of synonymous codon usage (SCU) in various genomes at inter and intra levels. Based on the codon usage and relative SCU results, five most frequently used amino acids were Leu, Ile, Phe, Met and Asn (Fig. 5). The most prevalent codons for the corresponding amino acids were UUA, AUU, UUU, AUA and AAU with the usage proportion values ranging from 45.53% to 48.95% (Supplementary Figure S2B). Cys was found to be the least used amino acid as detected in other selected Bombycoids. Biasness in SCU has been observed in many other lepidopterans to avoid errors due to the mis-incorporation of amino acids which depends upon various factors like nature selection (gene length and function), mutation bias (base mutation and GC content), etc. (12, 23). This A+T biasness may result in the change in amino acid composition similar to the other lepidopterans. Therefore, the codon usage analysis will have great significance in studying gene expression and evolutionary studies in silkworms.

**Figure 5.**
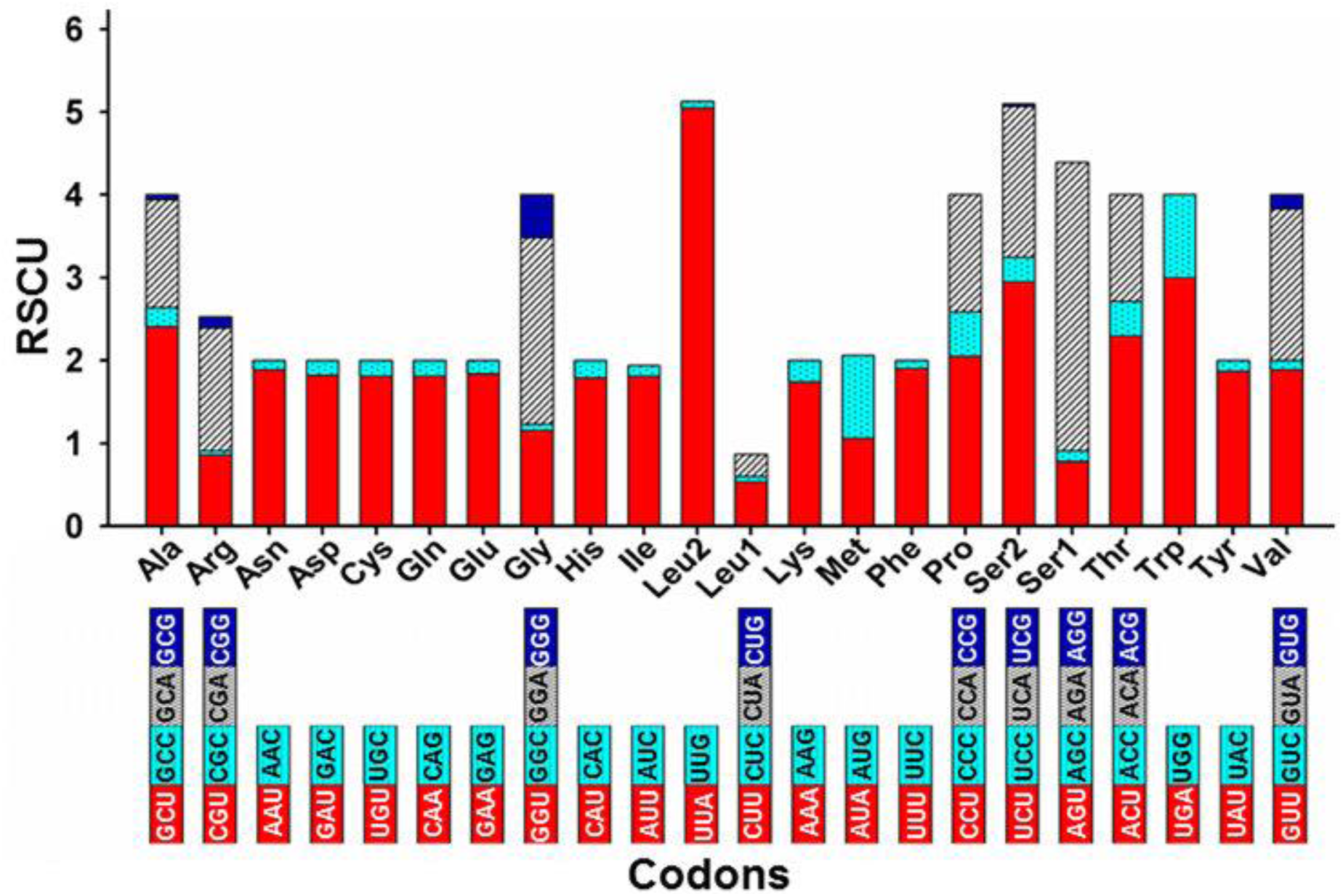
Relative synonymous codon usage (RSCU) of protein coding genes of *A. assamensis*. Termination codons were excluded in the study. Codon families are provided on X-axis and RSCU values on Y axis.

### Comparative phylogenetic analysis

Two phylogenetic trees were constructed based on 13 concatenated PCGs sequences (Fig. 6) and whole mitogenome sequences respectively (Supplementary Figure S6). Among various substitution models, GTR+I+G model was selected as the optimal model using AICc and BIC in jModeltest 2.1.7 software (46). The tree clustering showed that *A. assamensis* is a closely related member of Bombycoidea superfamily (Saturniidae+Bombycidae+Sphingidae) within the order Lepidoptera. It belongs to the Saturniinae subfamily of Saturniidae family. Within the subfamily, the tribes Saturniini (*A. assamensis*, *A. yamamai, A. pernyi, A. frithi, A. selene, S. boisduvalii, E. pyretorum*) and Attacini (*A. atlas, S. ricini*) formed sister groups.

**Figure 6.**
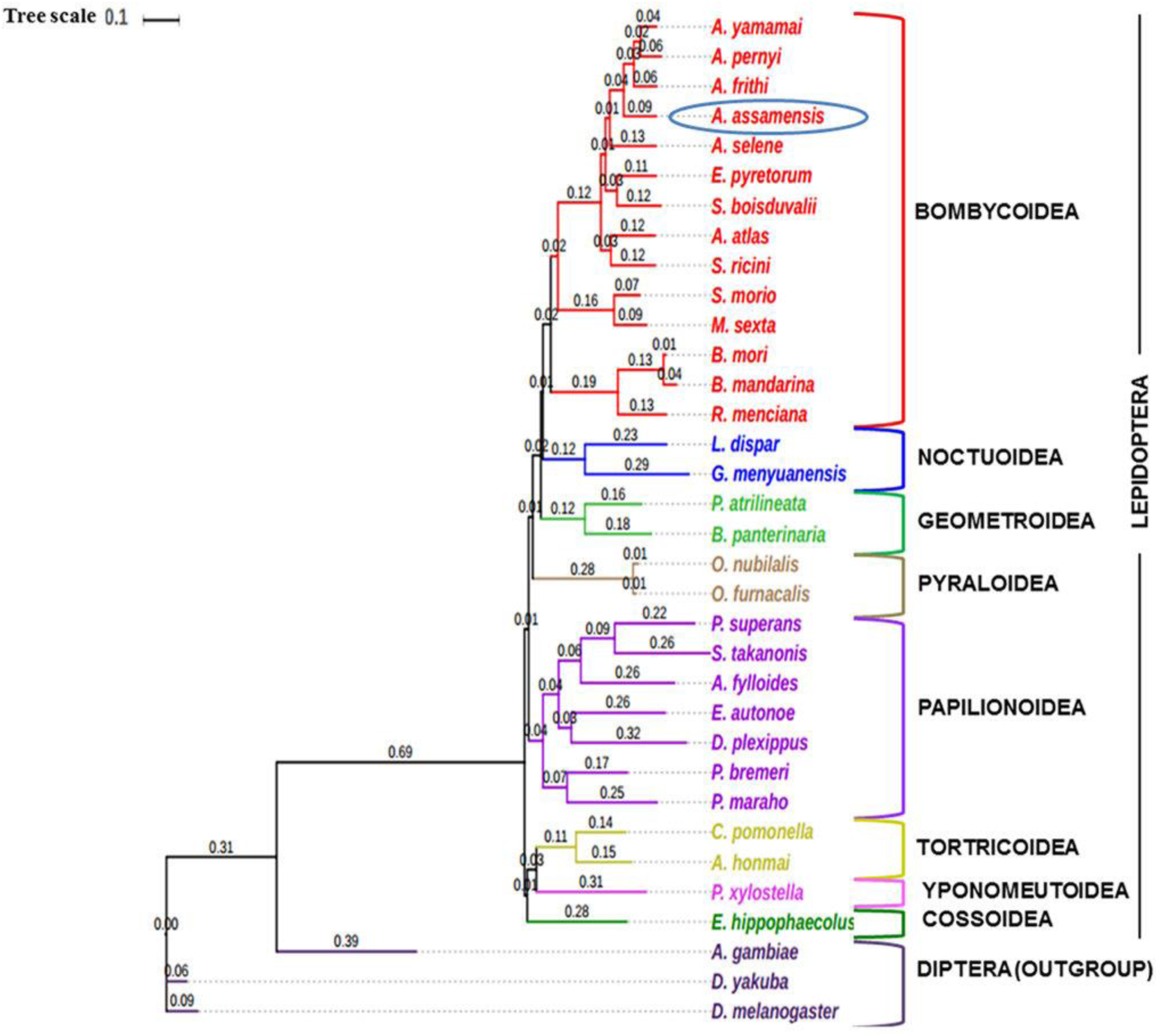
Molecular phylogeny inferred from concatenated nucleotide sequences of 13 protein coding genes (PCGs) of 34 organisms using Maximum Likelihood method. The scale for genetic distance is shown at the top of the figure and the numbers denote branch lengths of the tree.

Based on the PCGs dataset, Bombycoidea was found to be closest with Noctuoidea which further formed sister group with Geometroidea followed by Pyraloidea, Papilionoidea, Tortricoidea, Yponomeutoidea and Cossoidea. The closest sister cluster between the Bombycoidea and Noctuoidea was consistent with the previously reported studies (15, 23, 32). The sister group relationship between Noctuoidea and Geometroidea was observed using both datasets. However, more mitogenome sequence information on Bombycoidea superfamily is needed to gain deep insights into the families/superfamilies relationships within Lepidoptera. The present study support the fact that intra and inter family level relationships are well resolved within the superfamily Bombycoidea providing a better evolutionary relationship among the silk producing insects.

In summary, our study deciphered the complete mitogenome of *A. assamensis* which shares similarity with majority of the lepidopterans, particularly Saturniids, in several characteristics such as genome organization and content, PCG size and structure, AT/GC skewness, tRNA structure and anti-codons, OS and IGS regions. Our analysis indicates that *A. assamensis* PCGs evolved under strong purifying selection and tRNAs genes showed high base substitution/mismatches. In addition, higher nucleotide substitution (Transition+Transversion) in PCGs suggests high sequence divergence with respect to *B. mori* than with *M. sexta* and *A. pernyi.* A six-bp indel (deletion) in control region may have potential application as a marker for delineating other closely related *Antheraea sp.* and subsequent usage in phylogenetic studies at lower taxonomic levels. The largest *tRNA*^*Gln*^*-nad2* spacer found in the *A. assamensis* mitogenome may serve as another origin of replication. Our study assigns the taxonomic status of *A. assamensis* using optimal model based phylogenetic tree construction. This is the 12^th^ representative organism of Saturniidae family with a completely sequenced mitogenome. As *A. assamensis* is the sole producer of unique golden muga silk which is the backbone of Assam’s sericulture industry, our study on its mitogenomic landscape is an important addition to the existing genome informatics resources on silkworms.

## Materials and methods

### Sample processing, sequencing and assembly

The organism *A. assamensis* was reared in the experimental field of Central Muga Eri Research and Training Institute, Lahdoigarh, Jorhat, Assam, India. The leaves of *Persea bombycina* were used to feed the larvae following recommended package of practices (47). The fifth instar larval stage of organism was used for sample preparation for the present study with Sample ID-CMERI-Aa- 001. The sample was prepared in sterile condition by dissecting and chopping of larvae which was washed with 70% ethanol in order to avoid external contamination. The adequate amount of sample was utilized for mitogenome studies and remaining sample was stored in 95% absolute ethanol at -80°C freezer for future use. The total DNA was extracted using CTAB (Cetyl trimethylammonium bromide) based buffer and silica column. Subsequently, mitochondrial DNA was enriched from total DNA extracted once the integrity, quantity and purity of extracted DNA was confirmed through agarose gel electrophoresis, light absorbance and fluorescence method. The complete overview of sequencing and analysis of mitochondrial genome of *A. assamensis* is represented in Fig. 7.

**Figure 7.**
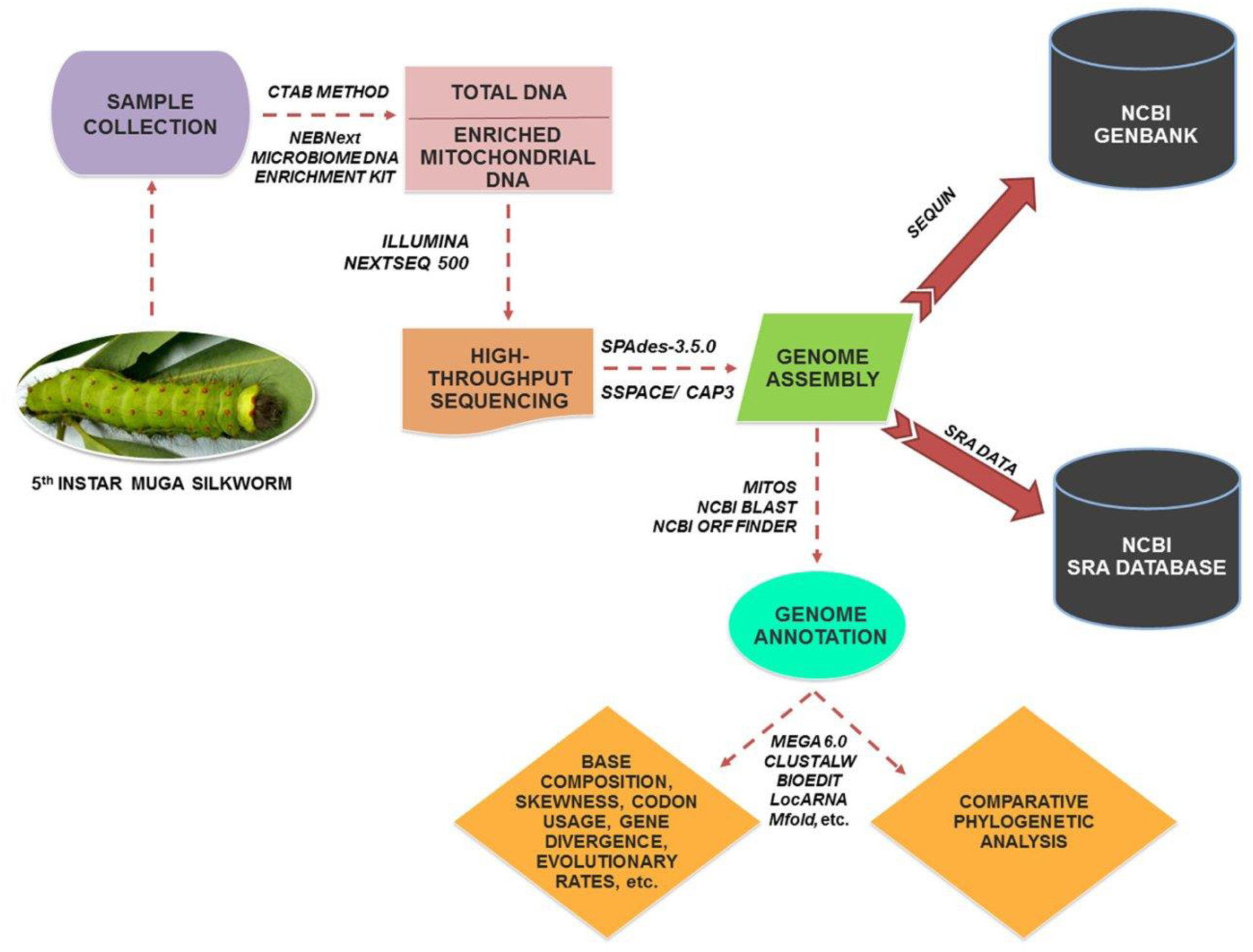
A Pictorial overview of the methodologies used for sequencing and analysis of *A. assamensis* mitogenome.

In the first step, preferential enrichment of mitochondrial DNA was carried out using NEBNext microbiome DNA enrichment kit (New England Biolabs, USA) which selectively removes eukaryotic nuclear DNA as it exist in CpG-methylated form unlike the mitochondrial DNA. In the next step, the enriched sample was used for mitochondrial DNA sequencing through the following methodology at Genotypic Technology Pvt. Ltd. Bengaluru, India. The enriched mitochondrial DNA was acoustically sheared to 300-500 bp using a specially designed Covaris microTube (for focused ultra-sonication). The fragmented mitochondrial DNA was cleaned up using HighPrep beads (MagBio Genomics, Inc, Gaithersburg, Maryland) for the removal of salts, primers, primer-dimers, dNTPs, etc. Further, fragmented DNA was subjected to end-repair, A-tailing and ligation with multiplex adaptors using the NEXTFlex DNA Sequencing kit (Catalogue # 5140–02, Bioo Scientific). The ligated DNA was cleaned with HighPrep beads and subjected to amplification via PCR as follows: initial denaturation at 98˚C for 2 min; 10 cycles of denaturation at 98˚C for 30sec, annealing at 65˚C for 30sec and extension at 72˚C for 60sec; and a final extension at 72˚C for 4 min) using the primers provided by NEXTFlex DNA Sequencing kit. Finally, the PCR product was purified with HighPrep beads, quantified and fragment range was assessed using Qubit flurometer and the Agilent D1000 Tape (Agilent Technologies) respectively in order to sequence on Illumina NextSeq500 (Illumina Inc, Sandiego, USA) through 2 × 150 bp paired-end chemistry. The obtained raw paired end data was de-multiplexed using Bcl2fastQ assessed with FastQC tool (48) to remove low quality bases (Q<30) and preprocessed using ABLT-Scripts (Genotypic technology, Bangalore India). The SPAdes-3.5.0 was used for sequence assembly preparation followed by the filling of gaps. Finally, scaffolding of assembled contigs and clustering were carried out with SSPACE and CAP3 programs respectively (49, 50).

### Genome annotation, visualization and comparative analysis

The assembled scaffolds were annotated using MITOS WebServer (51) which is widely used for the annotation of metazoan mitochondrial genomes due to its advanced annotation methodology. In the first step, the protein coding regions/genes were identified from the whole mitogenome sequence and confirmed using the NCBI (National Center for Biotechnology Information) ORF Finder (http://www.ncbi.nlm.nih.gov/gorf/gorf.html) by specifying the invertebrate mitochondrial genetic code. The initiation and termination codons were determined using NCBI BLAST, BioEdit and Clustal Omega by comparing the sequences of *A. assamensis* with the respective sequences published for other lepidopteran insects (52, 53). In the next step, the secondary structures of tRNAs were predicted using MITOS Server and in case of rRNAs, structure prediction was carried out by Mfold Web Server (http://unafold.rna.albany.edu/?q=mfold). Further, the number of overlapping or spacer regions between the genes was visualized and calculated manually. The control region was validated by comparing with the available sequences in GenBank and the tandem repeats in this region were determined through Tandem Repeat Finder (54). Then, the whole mito-map was constructed and visualized using Blast Ring Image Generator (BRIG) tool (55). Finally, the complete annotated file of mitochondrial genome was prepared using NCBI Sequin tool (http://www.ncbi.nlm.nih.gov/Sequin/) and the sequin file along with SRA data were submitted to NCBI GenBank.

The organism of present study, *A. assamensis,* was compared with the lepidopterans belonging to various levels of scientific classification (Table 1) for mitochondrial genome analysis. For this, the sequences of whole mitogenome, coding regions, tRNAs, rRNAs and control region were retrieved from the NCBI GenBank database.

### Comparative mitogenome analysis

Comparative mitogenome analysis was carried out in order to find out similarities and differences of *A. assamensis* with other insects in terms of characteristics like genome size, organization, structure, gene content, rearrangement and sequence similarity.

### Comparative analysis among protein coding genes (PCGs)

The PCGs of *A. assamensis* were compared with selected organisms based on their number, length, initiation codons and termination codons. The sequence similarity among the PCGs was determined by aligning the sequences of *A. assamensis* with selected Bombycoids using Clustal Omega. MEGA 6.0 tool was used for determining the gene-by-gene divergences in 13 PCGs in terms of phylogenetically informative sites, conserved sites as well as variable sites with some insects belonging to different classification levels (56). The same tool was used for estimating biasness in transitions and transversions of PCGs in order to determine nucleotide substitution patterns. Also, it was used to estimate Ka/Ks ratio of each PCG for determining the evolutionary rates. Further, the correlation between GC content and Ka/Ks ratio was studied in order to predict the effect of GC content on the evolutionary rates of PCGs.

### Comparative analysis among transfer RNAs (tRNAs)

The length, arrangement, secondary structures, variation in the secondary structures, etc. were studied among the selected organisms. Homologous sites in their secondary structures were identified by aligning the tRNAs sequences using LocARNA tool and the type/number of mismatches were calculated (57).

### Comparative analysis among ribosomal RNAs (rRNAs)

The number, type, location and length of rRNAs in *A. assamensis* were compared with the selected organisms in order to study the conservation pattern. In addition, sequence homology of rRNAs of *A. assamensis* with the selected organisms was determined; and gene-by-gene divergences among the rRNAs were studied using MEGA 6.0 tool.

### Comparative analysis among control region (A+T rich region)

The control region of *A. assamensis* was compared with selected organisms based on the length and location. Multiple sequence alignment (MSA) of control region was then carried out using Clustal omega and their conserved regions, repeats, indels, etc. were visualized using BioEdit tool (52).

### Comparative analysis among overlapping sequence (OS) and intergenic spacer (IGS) regions

The OS and IGS regions of *A. assamensis* mitogenome were compared with selected organisms (Table 1) in terms of number, length and location. The sequence homology of OS and IGS was determined through sequence alignment using Clustal Omega and BioEdit tools to find conserved motifs.

### Comparative analysis with respect to nucleotide composition, skewness and codon usage

The nucleotide composition of sequences of whole mitochondrial genome, concatenated and individual PCGs, tRNAs, rRNAs, spacers and control region was calculated using MEGA 6.0 software (56). The base composition skewness was also calculated for all the regions of mitogenome using the formula (E1 and E2) described by Junqueira et al 2004 (58).

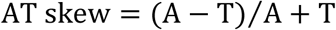

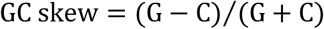

Where A, T, G and C denote the frequencies of respective bases.

The codon usage and relative synonymous codon usage (RSCU) values of PCGs were determined using MEGA 6.0.

### Comparative phylogenetic analysis

About 34 organisms representing 13 different families belonging to 8 super families within the order Lepidoptera were considered for phylogenetic analysis along with three different organisms belonging to the order Diptera as listed in Table 1. The phylogenetic relationship of *A. assamensis* was elucidated using two criteria’s: (i) concatenated nucleotide sequences of 13 PCGs and (ii) whole mitogenome sequences.

The nucleotide sequences of each PCG were translated via amino acid alignment using MAFFT algorithm in TranslatorX server which were then back translated into nucleotide alignment with GBlocks (59). The alignment of whole mitogenome nucleotide sequences was carried out in Clustal Omega. The statistical selection of best-fit models of nucleotide substitution was performed using jModeltest 2.1.7 based on PHYML 3.0. Maximum likelihood (ML) analysis of the sequences for each criterion was implemented based on corrected Akaike Information Criterion and Bayesian Information Criterion (46). The visualization of tree topology was carried out using iTOL v3 (Interactive tree of life) tool (60).

## Acknowledgements

The authors thank the Department of Biotechnology, Govt. of India, New Delhi for supporting the research through the UXCEL project (Sanction Order No: BT/411/NE/U-Excel/2013 dated 06.02.2014). The authors also express gratitude to MHRD and IITG for financial support in the form of fellowship.

## Author Contributions

DS wrote the main manuscript text and carried out data analysis; DK, HC and PS helped in data analysis; KN helped in the rearing and identification of the specimen used under the study; UB directed this study, reviewed analysis and revised the manuscript along with other authors.

